# Genomic evidence of genuine wild versus admixed olive populations evolving in the same natural environments in western Mediterranean Basin

**DOI:** 10.1101/2023.04.18.537296

**Authors:** Lison Zunino, Philippe Cubry, Gautier Sarah, Pierre Mournet, Ahmed El Bakkali, Laila Aqbouch, Evelyne Costes, Bouchaib Khadari

## Abstract

Admixtures between wild animals and plants and their domesticated relatives are widely documented. This can have positive or negative impacts on the evolution of admixed populations in natural environments, yet the phenomenon is still misunderstood in long-lived woody species, contrary to short-lived crops. Wild olive *Olea europaea* L. occurs in the same eco-geographical range as domesticated olive, i.e. the Mediterranean Basin (MB). Moreover, it is an allogamous and anemophilous species whose seeds are disseminated by birds, i.e. factors that drive gene flow between crops and their wild relatives. Here we investigated the genetic structure of western MB wild olive populations in natural environments assuming a homogenous gene pool with limited impact of cultivated alleles, as previously suggested. We used a target sequencing method based on annotated genes from the Farga reference genome to analyze 27 western MB olive tree populations sampled in natural environments in France, Spain and Morocco. We also target sequenced cultivated olive tree accessions from the Worldwide Olive Germplasm Bank of Marrakech and Porquerolles and from an eastern MB wild olive tree population. We combined PCA, sNMF, pairwise Fst and TreeMix and clearly identified genuine wild olive trees throughout their natural distribution range along a north-south gradient including, for the first time, in southern France. However, contrary to our assumption, we highlighted more admixed than genuine wild olive trees. Our results raise questions regarding the admixed population evolution pattern in this environment, which might be facilitated by crop-to-wild gene flow.

## 1 Introduction

Gene flows between domesticated species and their wild relatives have been identified in several studies in animals (Anderson et al., 2021; Johnsson et al., 2016; Mary et al., 2022) and plant species (Hauser & Shim, 2007; Le Corre et al., 2020). This phenomenon is noted when cultivated genomic variants occur in unmanaged naturally occurring populations in natural environments. These admixed populations raise the question of the impact of gene flow on the evolution of natural populations. For instance, the introgression of new genetic diversity inside wild genomes can accelerate their evolution by increasing the frequency of favorable alleles, but can also be detrimental resulting in outbreeding depression, i.e. a loss of fitness in hybrids compared to their parents (Ellstrand et al., 1999; Gering et al., 2019; Keller et al., 2000).

Cultivated to wild introgressions variants may be found in several plant species. Crop-to-wild gene flow occurs in maize and teosinte, its closest relatives, for instance, which has been found to lead to the acquisition of herbicide resistance in teosinte and, consequently, to high frequency of teosinte forms in maize fields (Le Corre et al., 2020). In some perennial species such as apple, major introgressions with the spread of alleles from the cultivated gene pool to wild populations in Europe have been documented (Cornille et al., 2019). The resulting admixed populations showed higher fitness than wild apple trees (Cornille et al., 2019; Feurtey et al., 2017). Gene flow from domesticated relatives has also been reported in natural chestnut and poplar populations. Admixed poplar populations have been found in France (Chenault et al., 2011) and along the Danube River in natural environment (Jelić et al., 2015), while the same scenario has been observed in chestnuts in Japan (Nishio et al., 2021). All these studies have proposed conservation measures for *in-situ* and *ex-situ* preservation of genuine wild populations by limiting gene flow, by replanting genuine wild genotypes far from domesticated forms or by protecting the connections of wild metapopulations which can breed and thereby protect themselves from random genetic deterioration (Feurtey et al., 2017; Jelić et al., 2015). The evolutionary consequences of crop-to-wild gene flows in the natural environment and on wild populations are still misunderstood, especially in perennial species.

Olive tree, (*Olea europaea* L.) is an iconic perennial species from the Mediterranean Basin (MB) which can live thousands of years. Cultivated (*Olea europaea* var. *europaea*) and wild (*Olea europaea* var. *sylvestris*) forms coexist within the same Mediterranean distribution range (Rubio de Casas et al., 2006; Zohary et al., 2016). Wild olive trees have an ancient evolutionary history in the MB (Besnard, Khadari, et al., 2013) indicated that three plastid lineages with a probable common ancestor dating from the Middle to Upper Pleistocene had diversified long before the Last Glaciation Maximum (26,500 to 19,000 BP (Clark et al., 2009)). They were subsequently impacted by glaciation, while some wild populations persisted in refugia (Besnard, Khadari, et al., 2013; Carrión et al., 2010). Olive lineages have been isolated in two distant areas, which could explain the current population genetic structure profile of wild olive trees. According to previous genetic studies, two main gene pools are identified, one in the eastern and another in the western/central MB (Besnard, Bakkali, et al., 2013; Besnard et al., 2001; Besnard, Khadari, et al., 2013; Diez et al., 2015). This eastern/western genetic differentiation is also found in other plants in the MB (Arroyo-García et al., 2006; Rodríguez-Sánchez et al., 2009). Cultivated olive trees emerged with the domestication of olive trees around 6,000 years BP (Kaniewski et al., 2012; Newton et al., 2014; Zohary et al., 2016). It is generally considered that the 4 center of primary olive domestication, from wild progenitors, is located in the Middle East, near the border between Turkey and Syria (Besnard, Khadari, et al., 2013; Kaniewski et al., 2012; Zohary et al., 2016).

It is currently impossible to distinguish between genuine wild, admixed and cultivated olive trees in the natural environment because of the absence of distinguish morphological features, so these forms can only distinguished using genetic markers (Besnard, Bakkali, et al., 2013; Diez et al., 2015; Gros-Balthazard et al., 2019; Julca et al., 2020; Khadari & El Bakkali, 2018; Lumaret & Ouazzani, 2001). The genetic diversity of cultivated olive trees is close to eastern MB wild olive trees (Besnard et al., 2001; Gros-Balthazard et al., 2019; Lumaret & Ouazzani, 2001), hence making it difficult to study gene flow between eastern MB wild and cultivated accessions. Conversely, in western MB, the genetic diversity of wild olive trees growing in natural areas is clearly different from cultivated accessions (Besnard, Bakkali, et al., 2013; Besnard, Khadari, et al., 2013; Khadari & El Bakkali, 2018), thereby enabling the identification of genuine wild olive trees as previously reported by Lumaret & Ouazzani (2001) using allozyme markers. Genetic pattern observed in naturally occurring populations was few impacted by crop-to-wild gene flow (Besnard, Bakkali, et al., 2013; Gros-Balthazard et al., 2019; Khadari & El Bakkali, 2018). The well-known genetic differentiation makes it a relevant model for investigating the genetic structure of populations in their natural environments and to infer potential gene flow between cultivated and wild olive.

Here, we investigated the genetic diversity of naturally occurring olive tree populations in the western MB. We assumed that the genetic pool of wild olive tree in the western Mediterranean area has not been impacted by introgressions from domesticated forms. We addressed the following questions: (1) What is the genetic structure of spontaneous olive trees in the western MB? (2) Are there genuine wild olive populations in this range? (3) Is there crop-to-wild gene flow in this region? We analyze genome-wide SNPs in olive trees from 27 natural sites ranging from southern France, Spain and Morocco. We included DNA from wild trees previously sampled in southern Turkey for the purpose of comparing diversity in these populations with the genetic pattern in the eastern MB wild gene pool (Gros-Balthazard et al., 2019). We sought to identify crop-to-wild gene flow and patterns of admixtures using data of cultivated accessions from western MB from the Worldwide Olive Germplasm Bank of Marrakech (WOGBM) and Porquerolles obtained with the same sequencing strategy (El Bakkali et al., 2019; Khadari et al., 2019).

## 2 Methods

### 2.1 Sampling of wild olive trees on a north-south gradient in the western Mediterranean Basin

Sampling of 27 assumed wild olive tree sites was conducted along a north-south gradient from southern France to southern Morocco in 2021 and early 2022. Sites were selected via the Conservatoire Botanique National Simethis database (http://simethis.eu) for southern France and northern Spain whereas for Corsica, central and southern Spain and Morocco, the delineation of wild populations was based on plastid polymorphism as reported by Besnard, Bakkali, et al. (2013). In addition to information from the Simethis database, we used environmental criteria to limit sampling of admixed olive populations and disregard olive orchards, agricultural and urban environments (Copernicus Land Monitoring Service, 2020). This resulted in the selection of 27 sites (Table 1; Figure 1A). At each of them, 13 to 15 individuals were sampled, representing a total of 400 sampled wild olive trees. At each site and for each tree, leaves were collected and immediately dried in silica gel for subsequent DNA extractions.

**Figure 1.**
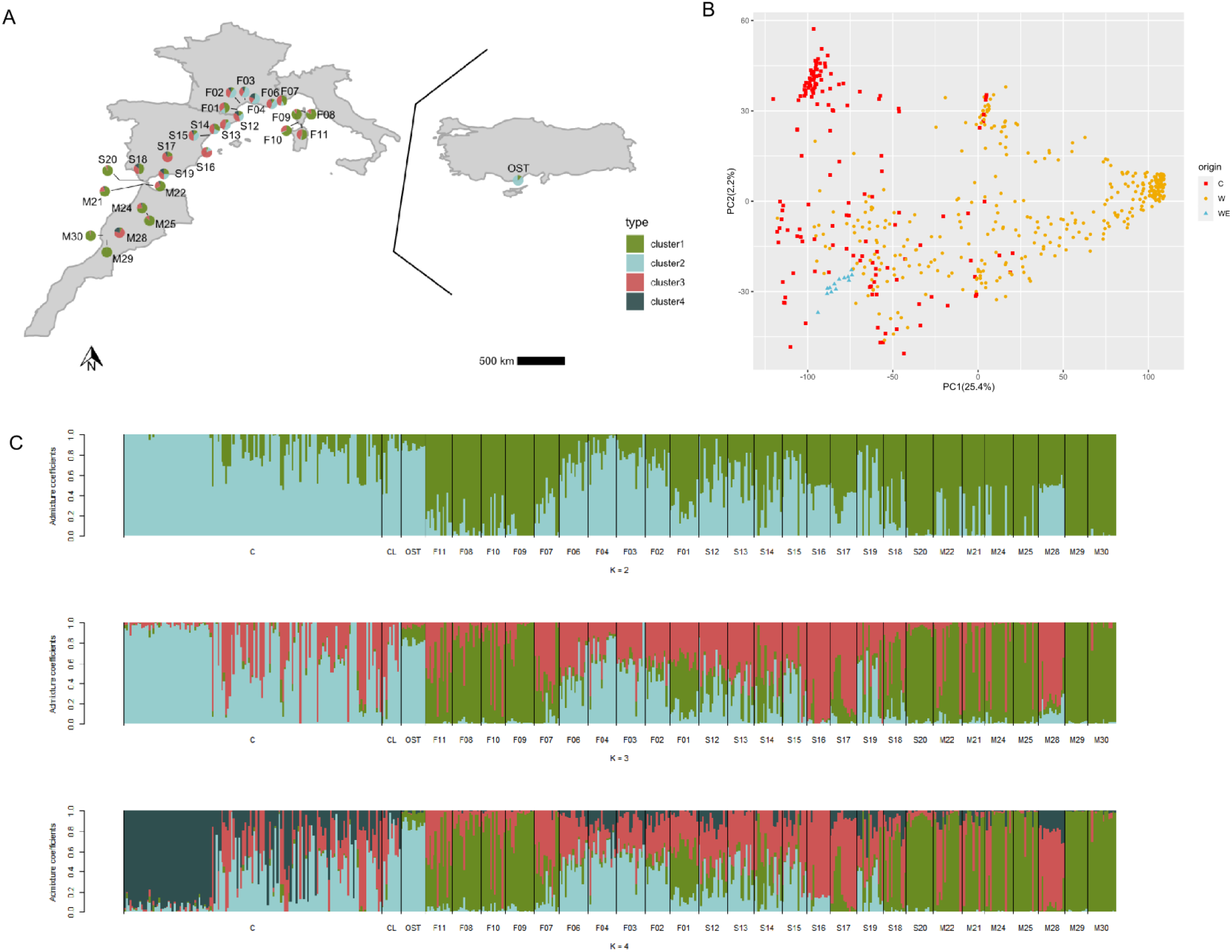
Result of the population structure analyses performed on the genome-wide SNPs diversity of natural populations of *O. europaea L.* collected in France (143), Spain (123), Morocco (96) and Turkey (13) and cultivated *O. europaea L.* from the western Mediterranean Basin (145), using 142,060 SNPs. (A) Geographical distribution of the populations and proportion of genetic cluster assignation. Pie chart at each location represents the fraction of individuals belonging to each genetic cluster as inferred by sNMF (K = 4). (B) PCA inferred with LEA. C: Cultivated, W: Western wild, WE: Eastern wild. (C) Genetic structure inferred by sNMF, each horizontal bar indicates individual assignment to a genetic cluster with K being the number of genetic clusters (K from 2 to 4).

**Table 1.**
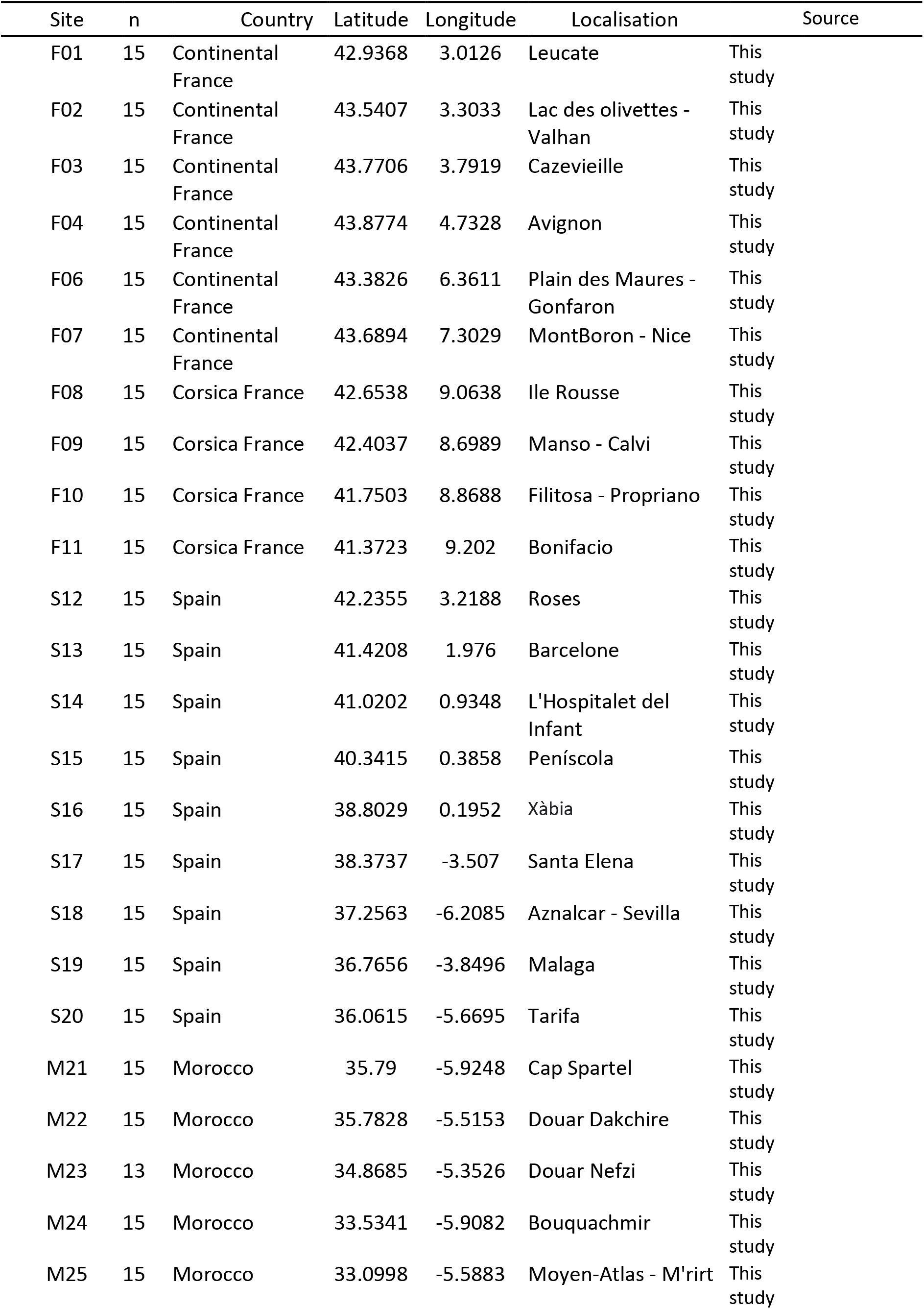
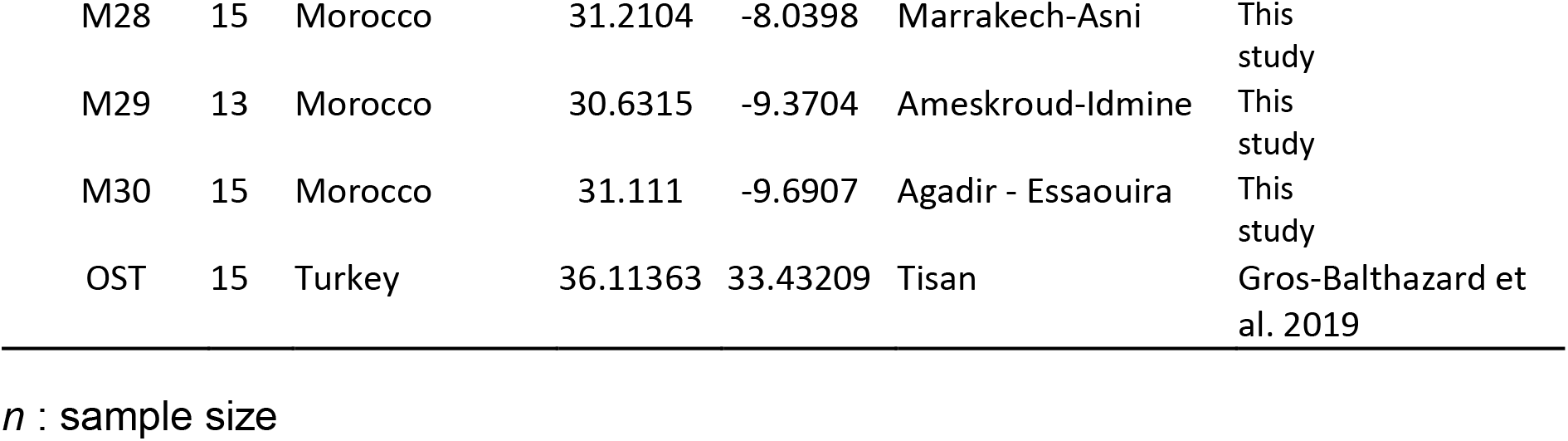
Summary of the localisation of the natural olive populations sampled for the study.

| Site | n | Country | Latitude | Longitude | Localisation | Source |
| --- | --- | --- | --- | --- | --- | --- |
| F01 | 15 | Continental France | 42.9368 | 3.0126 | Leucate | This study |
| F02 | 15 | Continental France | 43.5407 | 3.3033 | Lac des olivettes - Valhan | This study |
| F03 | 15 | Continental France | 43.7706 | 3.7919 | Cazevieille | This study |
| F04 | 15 | Continental France | 43.8774 | 4.7328 | Avignon | This study |
| F06 | 15 | Continental France | 43.3826 | 6.3611 | Plain des Maures - Gonfaron | This study |
| F07 | 15 | Continental France | 43.6894 | 7.3029 | MontBoron - Nice | This study |
| F08 | 15 | Corsica France | 42.6538 | 9.0638 | Ile Rousse | This study |
| F09 | 15 | Corsica France | 42.4037 | 8.6989 | Manso - Calvi | This study |
| F10 | 15 | Corsica France | 41.7503 | 8.8688 | Filitosa - Propriano | This study |
| F11 | 15 | Corsica France | 41.3723 | 9.202 | Bonifacio | This study |
| S12 | 15 | Spain | 42.2355 | 3.2188 | Roses | This study |
| S13 | 15 | Spain | 41.4208 | 1.976 | Barcelone | This study |
| S14 | 15 | Spain | 41.0202 | 0.9348 | L'Hospitalet del Infant | This study |
| S15 | 15 | Spain | 40.3415 | 0.3858 | Peníscola | This study |
| S16 | 15 | Spain | 38.8029 | 0.1952 | Xàbia | This study |
| S17 | 15 | Spain | 38.3737 | -3.507 | Santa Elena | This study |
| S18 | 15 | Spain | 37.2563 | -6.2085 | Aznalcar - Sevilla | This study |
| S19 | 15 | Spain | 36.7656 | -3.8496 | Malaga | This study |
| S20 | 15 | Spain | 36.0615 | -5.6695 | Tarifa | This study |
| M21 | 15 | Morocco | 35.79 | -5.9248 | Cap Spartel | This study |
| M22 | 15 | Morocco | 35.7828 | -5.5153 | Douar Dakchire | This study |
| M23 | 13 | Morocco | 34.8685 | -5.3526 | Douar Nefzi | This study |
| M24 | 15 | Morocco | 33.5341 | -5.9082 | Bouquachmir | This study |
| M25 | 15 | Morocco | 33.0998 | -5.5883 | Moyen-Atlas - M'rirt | This study |
| M28 | 15 | Morocco | 31.2104 | -8.0398 | Marrakech-Asni | This study |
| M29 | 13 | Morocco | 30.6315 | -9.3704 | Ameskroud-Idmine | This study |
| M30 | 15 | Morocco | 31.111 | -9.6907 | Agadir - Essaouira | This study |
| OST | 15 | Turkey | 36.11363 | 33.43209 | Tisan | Gros-Balthazard et al. 2019 |

$n$ : sample size

### 2.2 Reference set of cultivated and eastern wild accessions

In addition to the sample sites described above, leaves from 10 cultivated olive varieties were added to the sampling. These varieties were selected because of their significant presence in the French sampling area of natural populations (Khadari et al., 2019), which can be a potential source of introgression. Fifteen individual wild olive trees from Turkey in the eastern MB (Gros-Balthazard et al., 2019) were also added to create a genetically distinct group which will be considered as an outgroup (Figure 1A). Moreover, 135 cultivated varieties from the WOGBM, representative of the genetic diversity of olive resources in the western MB (El Bakkali et al., 2019) were considered as reference varieties to assess the introgressions from cultivated olive into wild populations. This last dataset was developed in a parallel study by our group that is focusing on cultivated olive (Supplementary Table S2). Overall, the experiment included 561 individuals.

**Table 2.** Summary information of genetic diversity for sampling sites of naturally occurring olive trees in the western and eastern Mediterranean Basin.

| | n | $H_o$ | $H_E$ | $F_{IS}$ |
| --- | --- | --- | --- | --- |
| *F01* | 15 | 0.142±0.195 | 0.141±0.178 | -0.003±0.263 |
| *F02* | 13 | 0.147±0.199 | 0.144±0.180 | -0.014±0.265 |
| *F03* | 15 | 0.145±0.213 | 0.130±0.177 | -0.086±0.234 |
| *F04* | 15 | 0.134±0.202 | 0.124±0.174 | -0.056±0.247 |
| *F06* | 15 | 0.148±0.202 | 0.142±0.180 | -0.032±0.249 |
| *F07* | 13 | 0.154±0.218 | 0.141±0.184 | -0.070±0.262 |
| *F08* | 15 | 0.127±0.197 | 0.121±0.173 | -0.029±0.262 |
| *F09* | 15 | 0.121±0.194 | 0.118±0.172 | -0.017±0.273 |
| *F10* | 13 | 0.135±0.205 | 0.126±0.176 | -0.053±0.271 |
| *F11* | 14 | 0.147±0.222 | 0.129±0.181 | -0.101±0.252 |
| *S12* | 15 | 0.158±0.214 | 0.146±0.184 | -0.057±0.242 |
| *S13* | 14 | 0.155±0.225 | 0.136±0.182 | -0.098±0.246 |
| *S14* | 15 | 0.154±0.204 | 0.149±0.184 | -0.026±0.254 |
| *S15* | 13 | 0.148±0.205 | 0.142±0.182 | -0.024±0.264 |
| *S16* | 12 | 0.187±0.274 | 0.144±0.187 | -0.206±0.319 |
| *S17* | 14 | 0.168±0.264 | 0.131±0.187 | -0.195±0.314 |
| *S18* | 12 | 0.147±0.204 | 0.141±0.182 | -0.028±0.271 |
| *S19* | 14 | 0.140±0.199 | 0.140±0.184 | 0.001±0.280 |
| *S20* | 14 | 0.123±0.201 | 0.115±0.174 | -0.043±0.269 |
| *M21* | 12 | 0.139±0.206 | 0.129±0.175 | -0.051±0.262 |
| *M22* | 15 | 0.141±0.196 | 0.135±0.173 | -0.030±0.246 |
| *M24* | 15 | 0.139±0.202 | 0.131±0.174 | -0.045±0.247 |
| *M25* | 13 | 0.127±0.197 | 0.123±0.174 | -0.020±0.270 |
| *M28* | 14 | 0.175±0.262 | 0.139±0.189 | -0.192±0.295 |
| *M29* | 12 | 0.114±0.198 | 0.111±0.176 | -0.018±0.307 |
| *M30* | 15 | 0.121±0.198 | 0.115±0.174 | -0.031±0.268 |
| *OST* | 13 | 0.117±0.183 | 0.116±0.166 | -0.007±0.271 |

n, number of genotypes; $H_E$ , expected heterozygosity; $H_o$ , observed heterozygosity; $F_{IS}$ , population level deviation from Hardy-Weinberg heterozygosity

### 2.3 Bait design

The cultivated *Olea europaea* var. *europaea* (cv. Farga) Oe9 genome assembly (Julca et al., 2020) was used as a reference to design target sequencing probes. This genome is 1.38 Gb. Baits were designed according to the following parameters: place 80 bp probes with 0.5x tilling targeting the first 640 bp of each of the 55,595 annotated genes available. For each gene, 1 to 4 baits were designed depending on its length. After quality filtration, this resulted in a total set of 210,367 baits representing 55,452 unique loci and a captured length of 16.8 Mb. The probes were designed and 7 synthesized by Daicel Arbor Biosciences, Ann Arbor, Michigan, USA. For this study, we only retained sequencing data targeted on assembled chromosomes. This subset represented 102,126 baits with a captured length of 8.2 Mb (Supplementary table S1).

### 2.4 Library preparation and sequencing

DNA was extracted from leaves using a mixed alkyl trimethylammonium bromide buffer (MATAB) and NucleoMag Plant Kit (Macherey-Nagel, Düren, Germany) as already described by Cormier et al. (2019) (Supplementary information). Individual genomic libraries for the NGS experiments were constructed with the NEBNext® Ultra™ II FS DNA Library Prep Kit (New England Biolabs, Ipswich, MA) with inputs ≤ 100 ng (Supplementary information). DNA was enzymatically sheared at an average 160 bp length before being tagged with the Unique Dual Index. Enrichment by capture was performed with biotinylated RNA probes (80 bp) as recommended by the provider using myBaits kits (Arbor Biosciences). A single dose of bait was used on a bulk of 48 normalized libraries. The sequencing was performed by MGX-Montpellier GenomiX on an Illumina® Novaseq^TM^ 6000 (Illumina Inc., San Diego, CA, USA) platform with an S4 flow cell. In addition to the target sequencing data set, we also sequenced four whole genomes (OES_E13_09, OES_F10_03, Picholine and Picholine Marocaine) to calculate the enrichment rate of the target sequencing method.

#### 2.4.1 SNP calling

Raw sequencing reads were first trimmed with FastP version 0.20.1 (Chen et al., 2018). The resulting data were then mapped on the reference genome Farga Oe9 genome assembly (Julca et al., 2020) using bwa-mem2 version 2.0 (Vasimuddin et al., 2019). The mapped reads were sorted with samtools version 1.10 (Li et al., 2009). Only primary alignment, properly paired and unique reads were kept. Duplicates were 8 removed using picard-tools version 2.24.0 (Broad Institute, 2019). From this clean alignment, GATK version 4.2.0.0 (Poplin et al., 2017) was used for the SNP calling according to GATK4 best practices (https://gatk.broadinstitute.org/hc/en-us/sections/360007226651-Best-Practices-Workflows).

#### 2.4.2 SNP filtering

After sequencing, we obtained 27,275,679 raw data. These were filtered with VCFtools version 0.1.16 (Danecek et al., 2011). The following filters were sequentially applied. With vcf-annotate, we first removed SNPs with a quality below 200 and clusters of 3 SNPs in 10 bases. VCFtools was used to remove indels, to only keep biallelic SNPs, to select SNPs with a minimum mean depth per site of 8, a minimum depth per site of 8 and a maximum mean depth per site of 400. Sites with >15% missing data were removed, then individuals with >20% missing data were also removed (Supplementary table S4). After filtration, 35 individuals were removed (1 cultivated, 2 eastern wild plants and 32 western wild plants). Only sampling sites with at least 12 individuals were considered in this study. M23 had only 6 individuals left and was therefore removed from the data-set. Filtering was carried out to exclude positions with fixed heterozygosity (>85%) and the final filtering was done to keep at least one minor allele count per site. After all filtering steps, we obtained 142,060 SNPs in the final data set, these were located on all chromosomes (Supplementary table S3).

### 2.5 Genomic analysis

#### 2.5.1 Genetic diversity and genetic structure

Genetic diversity measure was examined for each sampled sites, considered as distinct populations. We calculated diversity measures as expected heterozygosit (*H*_E_), observed heterozygosity (*H*_O_) and inbreeding coefficient (*F*_IS_) using Hierfstat version 05-11 (Goudet & Jombart, 2022).

Pairwise between population genetic differentiations were estimated with Weir and Cockeman fixation index (*F*_ST_) using the pairwise.WCfst function from the Hierfstat package version 05-11 (Goudet & Jombart, 2022). Support values were calculated per locus, for each pair of population, based on bootstraps procedure (Supplementary Table S5).

Genetic structure analyses were conducted using sNMF and Principal Component Analysis (PCA) from the LEA package version 3.11.3 (Frichot & François, 2015). Both analyses were performed on the dataset. It included 135 individuals from the WOGBM, originating from Spain, France and Morocco according to their passport data, 10 cultivated samples from southern France, 362 western MB wild individuals and 13 eastern MB wild samples. For sNMF, five repetitions per clusters (K) considered, were performed with K ranging from 1 to 10.

#### 2.5.2 Admixture assessment

Inference of the population history, with the admixture and split pattern were done using TreeMix version 1.13 (Pickrell & Pritchard, 2012). This software constructs admixture graphs using allele frequencies of current genetic populations to infer a graph of all ancestral populations related to a common ancestor. For this analysis, we grouped cultivated olives in four different genetic groups, depending on their sNMF assignment to genetic ancestral clusters: C0, C1, C3 and C4 (Supplementary information; Supplementary table S6). With these clusters, the 27 populations collected in western MB and the Turkish population, we did 100 TreeMix runs with a random SNP block size between 100 and 1000, from 1 to 10 migrations each when considering the M29 population as an outgroup. We inferred the optimum number of migrations with multiple linear models and the Evanno method implemented in the *OptM* package version 0.1.6 (Evanno et al., 2005; Fitak, 2021). TreeMix analysis was performed with 500 bootstrap replicates, which were used to build a consensus tree with Phylip version 3.697 (Felsenstein, 2005). We used BITE packages version 1.2.0008 (Milanesi et al., 2017) to display the trees.

## 3 Results

In this study, we analyzed 520 individuals, including 145 cultivars from Spain, France and Morocco, representing the MB olive diversity of cultivated olive trees (Supplementary table S2), a set of 362 wild trees from France, Spain and Morocco collected in 27 natural sites and 13 wild trees from southern Turkey.

### 3.1 Target sequencing efficiency

The average enrichment rate in the target sequencing experiment was 34 times higher than expected with whole-genome sequencing (Supplementary table S7). Moreover, for the bait on the chromosome annotated genes, 63.5% of the filtered SNPs (90,157) were on-target SNPs. The remaining was off-target SNPs. The on-target SNPs corresponded to sequences targeted by the baits. Conversely, the off-target corresponded to nonspecific and unintended sequences that can arise through sequencing (Supplementary table S7).

### 3.2 Genetic diversity and genetic structure

The average inbreeding coefficient (*F*_IS_) calculated on the different populations was on average 0. This is in accordance with the outcrossing mating system of olive tree. For 3 populations we detected *F*_IS_ values ranging from -0.086 to -0.101 (F03, F11, S13) 11 while 3 other ones *F*_IS_ values ranging from -0.192 to-0.206 (S16, S17 and M28) (Table 2). All of these populations might be resulting from admixture event.

The pairwise differentiation values between the studied populations (*F*_ST_) ranged from 0.01 to 0.42 (Figure 2). The Turkish eastern wild sampling site was the most genetically differentiated from the western MB sites, such as Corsica, F01 and F07 in Continental France, S14, S17, S18 and S20 in Spain, and all sites in Morocco (all above 0.2). Compared to the Turkish wild population, M29 from the southern limit of the olive distribution is the most differentiated population (*F*_ST_ = 0.42), while F04 from France was the least differentiated population (*F*_ST_ = 0.01). We revealed a high genetic differentiation between western MB wild populations and the eastern MB wild population

**Figure 2.**
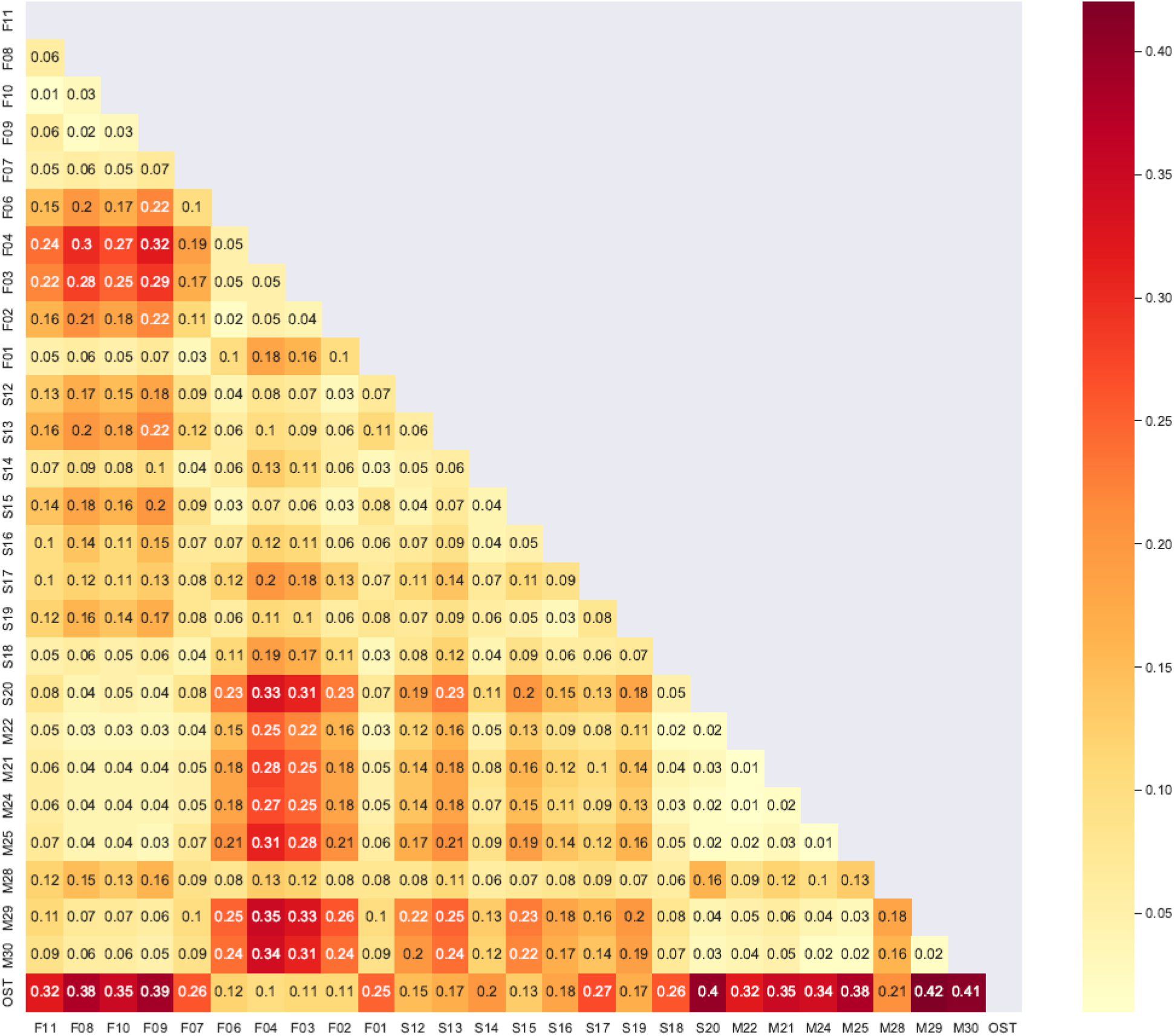
Heatmap of pairwise *F*_ST_ performed on the genome-wide SNPs diversity of 27 natural populations of *O. europaea* L. collected in western Mediterranean Basin in France (143), Spain (123), Morocco (96) and in the eastern Mediterranean Basin in Turkey (13).

In the principal component analysis (PCA), the first axis PC1 accounted for 25.4% of the variation and revealed an eastern-western genetic structure between western MB wild olive trees and eastern MB wild olive trees, with cultivated accessions mainly related to the eastern MB wild populations (Figures 1B and 3). On the first axis, we observed accessions from sampling sites in Corsica (F08 to F11, Figure 3B), S20, a large part of S18 from Spain (Figure 3C) and from all the sites in Morocco (Figure 3D), with the notable exception of M28, were clearly separated from the cultivated accessions and eastern MB wild accessions (Figure 3). All individuals collected in central southern France (F02, F03, F04 and F06; Figure 3A), one from Morocco (M28, Figure 3D) and some from north-central Spain (S12, S13, S15 and S16; Figure 3C) clustered with cultivated accessions as shown in the Figures 1B and 3 (left side of the PCA). This profile suggests admixture events. Several other individuals collected in eastern and western France (F01 and F07; Figure 3A), in Morocco (M21, M22, M24 and M25; Figure 3D) and in Spain (S14, S17, S18 and S19; Figure 3C) also exhibited a pattern of admixture with the cultivated accessions. The second axis, i.e. PC2, accounted for 2.2% of the observed variability, highlighting two subgroups within cultivated trees. The first subgroup includes cultivated genotypes mostly from Spanish varieties such as “Picual” and three Moroccan varieties including “Picholine Marocaine”. The other cultivated group included several varieties from Spain, Morocco and France (Figure 1B).

**Figure 3.**
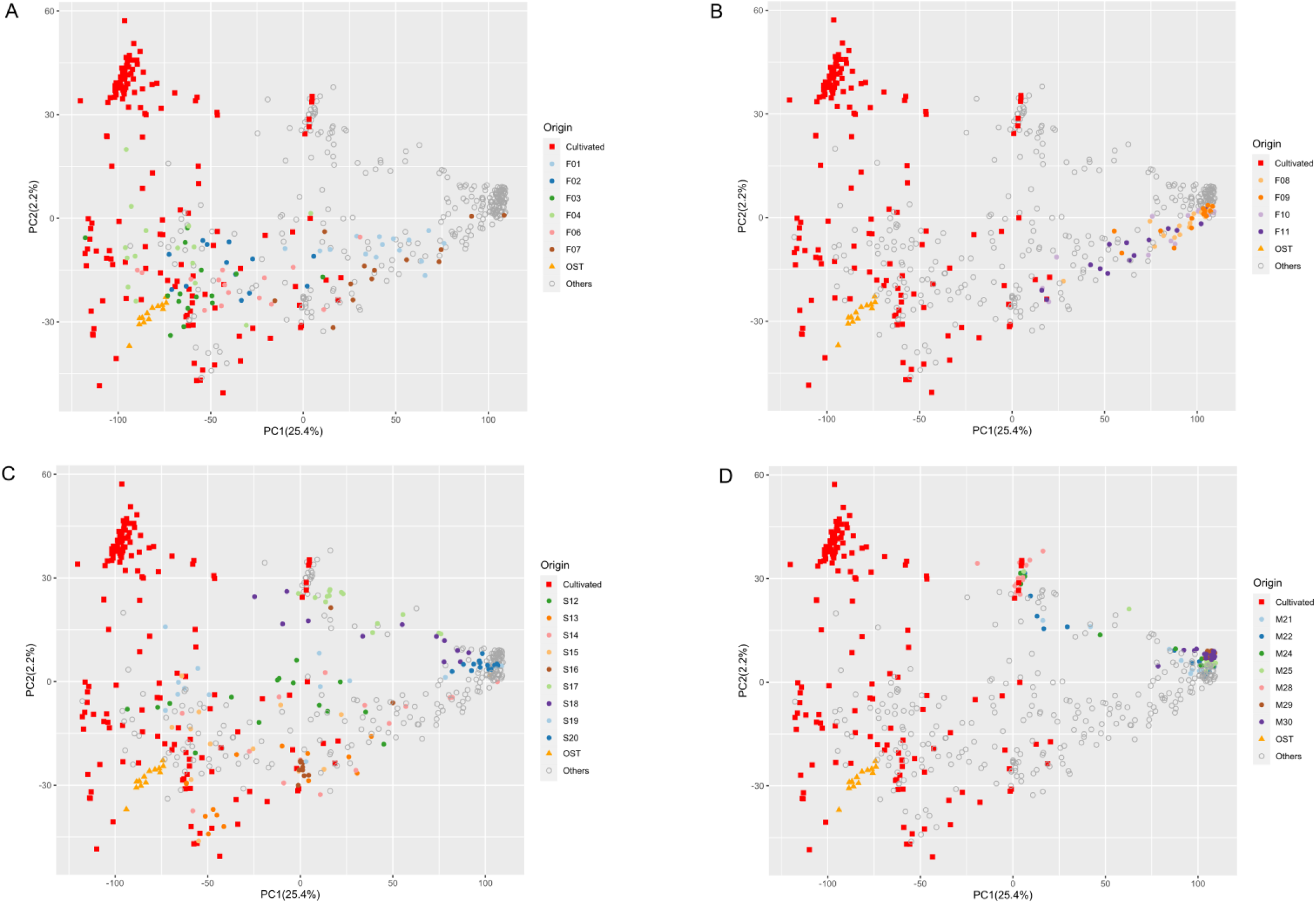
Detailed genetic structure of *O. europaea* L. populations using PCA analysis. Red boxes represent cultivated individuals, orange triangles represent Turkish population with the (A) French continental populations, (B) Corsican French populations, (C) Spain populations, (D) Moroccan populations.

A similar pattern was supported by the sNMF analyses. According to the cross-entropy criterion, only K from 2 to 4 were considered suitable to explain the western MB natural olive tree genetic pattern (Figure 1C; Supplementary figure S3). The wild Turkish olive population (OST) was assigned to a specific cluster from K=2 to K=4 (in blue) regardless of the admixture model examined, thereby supporting the existence of a structure between western MB and eastern MB natural populations. At K = 2, the cultivated and eastern MB olive trees collected in natural sites were assigned to cluster 1 (in blue) and the western MB wild natural olive trees in Morocco, Corsica, South Spain, France (F01 and F07 sites) were mainly assigned to cluster 2 (in green). Olive trees from the F02, F03, F04 and F06 sites in France were mainly assigned to cluster 1 (in blue). Trees collected from S12, S13, S15, S16, S17 S19 and M28 sites spread in the two clusters. At K = 3, these sampling sites were mostly assigned to cluster 3 (in red), particularly for individuals from populations M28, S17, and S16. With K = 4, a fourth cluster (in dark blue) was noted within the cultivated cluster, corresponding to the same first subgroup described in the PCA results above (Figures 1B and 3).

By combining three analyses (i.e. PCA, sNMF and pairwise *F*_ST_; Figures 1 and 3), particularly by considering the left and central part of the PCA (Figures 1B and 3) and the cluster 3 (in red; Figure 1C) from the sNMF analyses, we have several arguments strongly suggesting admixtures between natural olive trees and cultivated ones. It seems to be consistent with *F*_ST_ values, with much lower levels of differentiation between cultivated olive accessions and wild olive accessions (Figure 2). Accordingly, all olive trees mapped in the right side of the PCA (Figures 1B and 3) and assigned to the cluster (in green) regardless of the admixture model examined (Figure 1C) and considered as genuine wild.

### 3.3 Inference of population admixture and gene flow

The tree inferred by TreeMix was ranked using M29 population. This population was chosen because all of the individuals collected in this site belong to the same cluster, referred as the western MB wild cluster (in green; Figure 1C). This genuinely western MB wild olive also appeared to be the most genetically distinct from eastern MB wild olive (*F*_ST_ = 0.42) and from the cultivated accessions (Figure 1).

The TreeMix analysis revealed the highest divergence between M29 and C3 (cultivated) and OST (Turkish population). The lowest divergence (below 0.015) from M29 were found for with almost all the Moroccan sites (except for M28), with S20, F08, F09, F10 and F11. A second group of sampling sites was found with a divergence from M29 of 0.017 to 0.024, including S18, M28, F07, F01 and S17. Accessions from sampling sites in Spain, except for S20, had a genetic divergence of >0.032 from M29 and were closer to cultivated groups (<0.012 genetic divergence between S16 and C3). Accessions from the French F06, F02, F03 and F04 sites were found to be grouped with the cultivated clusters C3 and C0. TreeMix inferred a low divergence between C4, C1 and OST (around 0.005). Two gene flow events were inferred, with the first one being from cultivated and French populations from the center to M28, with a weight of (w = 0.425). The second one was from M22 to northern Spain sites (w = 0.485) (Figure 4).

**Figure 4.**
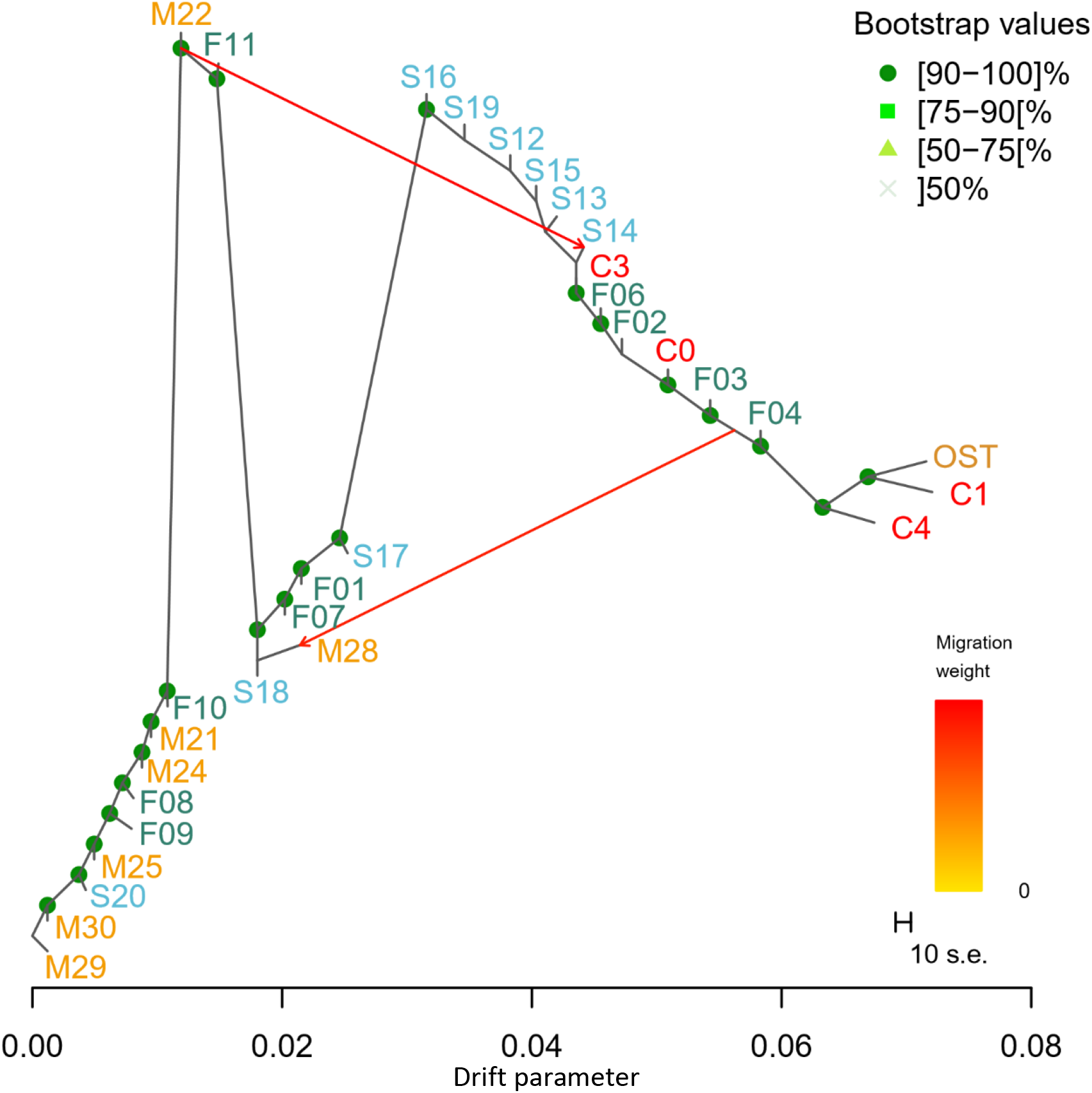
Tree inferred by TreeMix analysis on natural *O. europaea* L. populations from the western and eastern Mediterranean Basin and cultivated accessions. Genetic divergence is represented by the horizontal difference between populations. The vertical bars are only graphical representations and are not taken into account in the analysis. C0, C1, C3 and C4 are groups of cultivated accessions (Supplementary information).

## 4 Discussion

Wild olive molecular identification and characterization are important to assess the genetic diversity of this species, in addition, to evolutionary history and to investigate local adaptation, especially for wild relative crops such as olive, where wild olives and cultivated olives coexist in the same area (Rubio de Casas et al., 2006; Zohary et al., 2016). As crop-to-wild gene flows likely occur, deciphering their genetic relationships can help to understand the impact of crop on natural populations, their demographic histories and to explore new sources of genetic diversity. In this study, our aim was to identify genuine wild olive populations by investigating the genetic structure and diversity patterns of olive trees evolving in the natural environment in the western MB. We sampled allegedly wild olive trees over a large geographic area in the western MB, eastern MB genuine wild olive trees and cultivated olive trees. We analyzed this large panel of population using SNPs from target sequencing. This if the first time target sequencing has been used to study genomic variation in olive tree—it enabled a genome scan of many individuals while accessing more than 140,000 SNPs distributed throughout the genome. This represents a major advance over previous methods in similar studies using SSR-based molecular analyses (Besnard, Bakkali, et al., 2013; Khadari & El Bakkali, 2018) or SNPs analyses from RNAseq (Gros-Balthazard et al., 2019). We documented the presence of genuine wild olive tree populations in the western MB, in southern France, Spain and Morocco. However, contrary to our assumption, our analysis suggested that admixed populations are more frequent than genuine ones.

### 4.1 Genetic variation patterns in natural and cultivated olive trees highlight the persistence of genuine wild olive populations in the western Mediterranean Basin

Based on a large set of SNP markers, located all over the genome, we identify a very strong genetic differentiation between the natural populations collected in the western MB and those in the eastern MB. This confirmed the results of previous studies using microsatellite markers highlighted two distinct gene pools in the eastern and western MB. In these previous studies, the western/central MB gene pool clustered wild populations from Greece to Morocco (Besnard, Bakkali, et al., 2013; Khadari & El Bakkali, 2018) while the eastern MB gene pool clustered wild populations from Greece to the Levant. The eastern MB gene pool was represented here by one Turkish population (OST), i.e. a genuine wild population previously revealed using plastid DNA polymorphism (Besnard, Bakkali, et al., 2013; Besnard, Khadari, et al., 2013), SSR markers (Khadari & El Bakkali, 2018) and SNPs from RNAseq (Gros-Balthazard et al., 2019). Our results reflects the typical long evolutionary history of Mediterranean species as described by (Blondel et al., 2010).

Our study confirmed that the wild eastern MB olive population from Turkey is very closely related to cultivated olives, as shown by different approaches (PCA, sNMF, *F*_ST_ and TreeMix). This finding is consistent with the olive domestication history which was likely the results of genetic selection of eastern MB wild trees (Besnard, Khadari, et al., 2013; Gros-Balthazard et al., 2019; Khadari & El Bakkali, 2018). Moreover, we combined several approaches, i.e. PCA, sNMF, pairwise Fst estimation and inference of splits in mixture in populations and clearly identified a genetic group of olive trees strongly differentiated from the eastern wild and cultivated olive as previously found by Gros-Balthazard et al. (2019) which probably constitute genuine wild olive trees.

Finally, within cultivated accessions, we identified two genetic groups. The first one similar to OST and which might be composed of varieties likely issued from two processes, primary selection in the east and secondary diversification in the central and western Mediterranean areas as proposed by Khadari & El Bakkali (2018). The second group essentially consisted of Spanish and Moroccan varieties that were highly differentiated from OST and which might be composed of western MB cultivated olives through a secondary diversification process mainly via selection involving crossing between ancient varieties such as Gordal Sevillana and Lechin de Granada, as shown by Diez et al. (2015).

### 4.2 Wild versus admixed olive trees: an evolutionary history impacted by domestication

As discussed above, we identified genuine wild olive trees in the western Mediterranean basin in Spain, Corsica (France) and Morocco as already shown by Besnard, Bakkali, et al. (2013). The genetic diversity observed in Corsican and Moroccan populations were very close suggesting a common ancestral history for these populations. We also discovered, for the first time, the presence of genuine wild olive trees in continental France at Mont-Boron (eastern-south; F07) and near Leucate (western-south; F01). The abundance of wild olives trees in the western MB has been well characterized (Besnard et al., 2001; Besnard, Khadari, et al., 2013; Gros-Balthazard et al., 2019; Khadari & El Bakkali, 2018; Lumaret & Ouazzani, 2001). However, populations initially characterized as genuine wilds according to plastid DNA polymorphism and SSR markers, for instance, in Spain (Besnard, Bakkali, et al., 2013) were found admixed in our study. These sites were in habitats considered to be little or not at all impacted by human activities, e.g. in natural reserves, remote from urban centres or areas with olive orchards. Even within some sites, from a genetic viewpoint, several individuals were considered genetically to be wild, while others were very admixed with cultivated olives. This high genetic admixture intensity was unexpected in the western MB based on previous studies (Besnard, Bakkali, et al., 2013; Besnard, Khadari, et al., 2013; Besnard et al., 2018; Gros-Balthazard et al., 2019; Khadari & El Bakkali, 2018). Our findings have provided new insights into the evolutionary history of olive trees in natural western MB habitat.

### 4.3 What factors could influence the prevalence of admixed populations in natural environments?

We obtained clear evidence in this study on the substantial presence of admixed populations within the natural olive populations which were previously reported to be little impacted by crop-to-wild gene flow (Besnard, Bakkali, et al., 2013; Khadari & El Bakkali, 2018). This unexpected observation was unlikely the result from a sampling bias since we investigated olive trees from several sites already analyzed and considered as being genuine wild populations in previous studies (Besnard, Bakkali, et al., 2013; Besnard, Khadari, et al., 2013). Recent phylogenomic and population structure investigations revealed genetic admixtures during olive domestication thus highlighting the impact of domesticated alleles on two wild olive trees for the western MB (Julca et al., 2020). However, these authors analyzed very limited sampling of wild olive trees (7 olive trees from the western MB), while in our investigations, we analyzed 362 trees sampled from 27 natural sites in Spain, Morocco and France—this sampling covered the olive natural distribution range in the western MB. Through TreeMix analysis and comparing genomic variation comparison in these 362 olive trees to the genuine Turkish natural population and to cultivated olive trees, we, therefore, were able to depict gene flow between wild and cultivated olives (see Figure 4). The observed patterns highlighted the two kinds of gene flow events: from cultivated to wild 18 populations in Morocco, France and Spain and from wild to cultivated or already admixed populations in Spain.

These gene flows could have been driven by several factors. First, the mating system of olive, which, is allogamous, pollination mostly depends on wind (anemophilous pollination) and seed dissemination relies on birds (zoochorous dissemination). These forms of dissemination may occur over long geographic distances (>50 km) (Spennemann & Richard Allen, 2000). Second, cultivated and wild olive trees share the same climatic and ecological niches, the geographic proximity between them increases the possibility of gene flow and events of admixture (Besnard et al., 2018). Third, there could be cultivated versus wild olive tree pollen competition: monocultures and single-varietal olive orchards are responsible for broad dissemination of pollen from orchards (thousands of trees), whereas wild populations are often composed of few individuals. Wild olive pollen is thus less abundant. Fourth, gene flow between cultivated and wild olive trees may increase genetic diversity in admixed populations. Associated new variants or combination might be better adapted to the local environment, promoting an acceleration of local adaptation of a species to an environment is a recognized evolutionary force explaining the occurrence of admixed populations (Ellstrand et al., 1999). Evolutionary factors such as allogamy (Saumitou-Laprade et al., 2017) and cultivated versus wild olive tree pollen competition (see above) may not be sufficient to explain the large frequency of admixed populations versus genuine wild populations. Here we assume that admixed olive trees could have a better adaptive potential to their natural environment, as this has been previously demonstrated in several short-lived and annual crops (Feurtey et al., 2017; Johnsson et al., 2016; Le Corre et al., 2020). This assumption is supported by the findings of genetic investigations on natural olive trees in Australia (Besnard et al., 2014). These 19 authors hypothesized that hybridization between two introduced *Olea* species, *Olea europaea* subsp. *europaea* and *Olea europaea* subsp. *cuspidata*, overcame the lack of diversity after their introduction bottleneck, thereby facilitating their establishment. Strikingly, to our knowledge, this assumption has yet to be investigated in long-lived woody plants such as olive trees, even though knowledge on the impact of admixture on the evolution of natural populations could help guide appropriate conservation strategies in forest areas and other natural ecosystems.

### 4.4 Consequences of extensive hybridization of wild olive via domesticated olive introgression and conservation recommendations

The future of genuine wild genotypes might be threatened by the gene flow we highlighted here. For instance, extensive gene flow could ultimately lead to complete replacement of wild populations by admixed genotype (Ellstrand et al., 1999). However, in global change context, this gene flow could enhance adaptation to a changing environment. Our study offers new opportunities for more in-depth studies on this introgression process. We identified three different compartments, i.e. a genuine wild olive compartment, a cultivated one and an admixed one, that could be study to address this long-standing question. Conservation policies on wild olive trees should take into account the risk of introgression from cultivated alleles and by the impact of climate change. The naturally occurring olive trees sampled here were positioned on a north-south gradient with different environmental conditions, which could facilitate studies on their potential local adaptation to changing climatic conditions. Our study finding may provide a basis for designing new conservation measures to protect genuine wild genotypes, *in-situ* and *ex-situ*, including repositories of wild genetic diversity not impacted by artificial selection.

## 5 Conclusion

In this study we assessed the genetic structure of natural olive populations from the western Mediterranean Basin. We confirmed that the western MB genuine wild olive is genetically well differentiated from eastern MB wild olive as well as cultivated forms. We detected its presence in France, Spain and Morocco. We also found many admixed populations resulting from strong crop-to-wild gene flow. The presence of admixed olive populations in the same distribution area as genuine wild populations raises questions on the reasons for their predominance in the natural environment and on designing conservation strategies for both compartments. Finally, the two genetic patterns revealed by our investigations could be considered as a suitable model for investigating two core questions, the first on the admixture nature, i.e. what domesticated genomic alleles/regions would be suitable for introgression in wild genomes? The second question is related to local adaptation: are wild better locally adapted than admixed olive trees?

## Author Contributions

Bouchaib Khadari, Philippe Cubry and Evelyne Costes designed the research. Lison Zunino, Ahmed El Bakkali and Bouchaib Khadari collected the plant material. The DNA extraction and library preparation were done by Lison Zunino, Pierre Mournet and Laila Aqbouch. Gautier Sarah and Lison Zunino did the bioinformatic analysis. Lison Zunino, Philippe Cubry and Bouchaib Khadari performed the data analysis. Lison Zunino wrote the manuscript with the help of Bouchaib Khadari, Philippe Cubry and Evelyne Costes. All authors contributed to written work led by Lison Zunino. Bouchaib Khadari obtained the funding.

## Acknowledgements

We thank Guillaume Perez, Magalie Delalande, Karine Loridon, Juliette Arzul and Sylvia Lochon-Menseau for helping us in wild olive sampling from in situ areas; Anaïs Fossot, Hélène Vignes and Ronan Rivallan for the laboratory help. We acknowledge the helpful comments from Joëlle Ronfort and the English corrections from David Manley.

A part of a field work has been supported by the Conservatoire Botanique National Méditerranéen de Porquerolles, l’Institut Agro Montpellier and INRA Morocco. This work has been realized with the support of MESO@LR-Platform at the University of Montpellier, the financial support of CIRAD, and the technical support of the bioinformatics group of the UMR AGAP Institute, member of the French Institute of Bioinformatics (IFB) - South Green Bioinformatics Platform Lison Zunino received a PhD scholarship from the French government. This study was funded through Labex AGRO 2011-LABX-002, project ClimOliveMed n° 2003-001 (under I-Site Muse framework) coordinated by Agropolis Fondation.

## Data Accessibility and Benefit-Sharing Section

Raw sequence reads are deposited in the European Nucleotide Archive (PRJEB61410).

Snakemake workflow of the SNP calling available here: https://forgemia.inra.fr/gautier.sarah/ClimOlivMedCapture

**Benefits sharing**: Plant material was collected in three countries France, Spain and Morocco. Material sampling in France and Spain has been performed under supervision of the Conservatoire Botanique National Méditerranéen (Hyères France) and the International Olive Council (Madrid, Spain). A research collaboration was developed with one scientist from INRA Morocco who is included as co-author.

