## Supplementary information 1 for "Genomic evidence of genuine wild versus admixed olive populations evolving in the same natural environments in western Mediterranean Basin"

**Table of Contents:**

| **Supplementary Methods** | Page 2 |
| --- | --- |
| **Supplementary Figure S1** | Page 3 |
| **Supplementary Figure S2** | Page 4 |
| **Supplementary Figure S3** | Page 5 |
| **Supplementary Table S1** | Sep. File |
| **Supplementary Table S2** | Page 6 |
| **Supplementary Table S3** | Page 12 |
| **Supplementary Table S4** | Page 13 |
| **Supplementary Table S5** | Sep. File |
| **Supplementary Table S6** | Page 14 |
| **Supplementary Table S7** | Page 18 |

### Methods

#### DNA extraction:

DNA extraction was performed on leaves dried after a minimum of 10 days in silica gel. 25 mg of dried samples were sorted in a Deepwell 96 well plate (Corning 1.1mL Deepwell ThermoFisher Scientific) with beads. The plate was grinded using GenoGrinder (SPEX SamplePrep). The DNA was extracted using a MATAB buffer-based extraction method. Genomic DNA was quantified with Fluoroskan (Thermo fisher Scientific Inc). The quality was checked using agarose gel electrophoresis and Nanoquant plate (Tecan)

#### Library preparation:

The input DNA was fragmented for 10 minutes with fragmentase enzymes. The fragmentation size was checked with Tapestation with a mean average size of 160 bp. The adaptors were ligated then the samples were purified with AMPure beads (Beckman Coulter, Inc) at 0.8X to do a low and high size selection of 200 bp. Unique Dual Index (NEBNext Multiplex oligos for Illumina) were added to the adaptor-ligated DNA before doing an enrichment PCR as required for NovaSeq experiments. Libraries were amplified for 11 cycles and cleaned with AMPure beads at 0.9X. The library quality was checked with Tapestation. Each library was quantified by qPCR on a LightCycler (Roche Molecular Systems, Inc) before being mixed together on a 48 libraries equimolarity pool. Hybridization capture for targeted NGS protocol is applied on each pool. The baits are added in a pool before hybridizing with beads for at least 16 hours at 60°C. The beads are then washed and the Bait-target hybrids are amplified by PCR for 12 cycles. The reaction is purified with AMPure beads. The captured library quality was verified by Tapestation.

Each capture-pool was quantified by qPCR on LightCycler before being grouped in one final pool of 384 libraries based on equimolarity.

#### Cluster assignment to create subgroup inside cultivated:

Using sNMF from LEA packages V3.11.3 (Frichot and François 2015), we did structure analyses of the subgroup of cultivated (cultivated from west, including varieties from Spain, Morocco and France). Five runs were performed with K from 1 to 10. According to the cross-entropy, only K of 4 was represented in the results. Using this, we did an assignment of each variety to a cluster, if the variety was assigned to a cluster more than 70%. If the individual is not assigned to any cluster with these parameters, then it is assigned to an admixed cluster, C0. The C2 group was composed only with 3 individuals, which are not sufficient to a TreeMix analysis, we decided to put them in the C0 group and remove C2.

**Supplementary figures:**

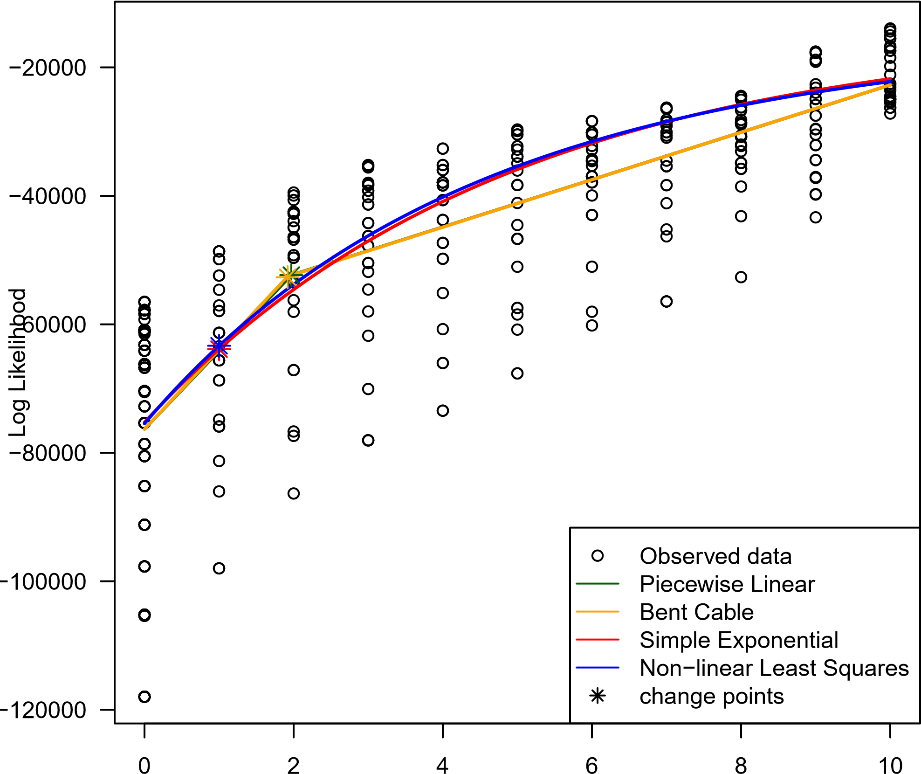

**Figure S1**. Multiple linear models of possible migration events between 27 naturally occurring populations, eastern wild Mediterranean population and cultivated accessions of olives using TreeMix. One hundred TreeMix runs were done with a random SNPs block size between 100 and 1000, from 1 to 10 migrations each with M29 as outgroup. The analysis was performed with *OptM* package.

m (migration edges)

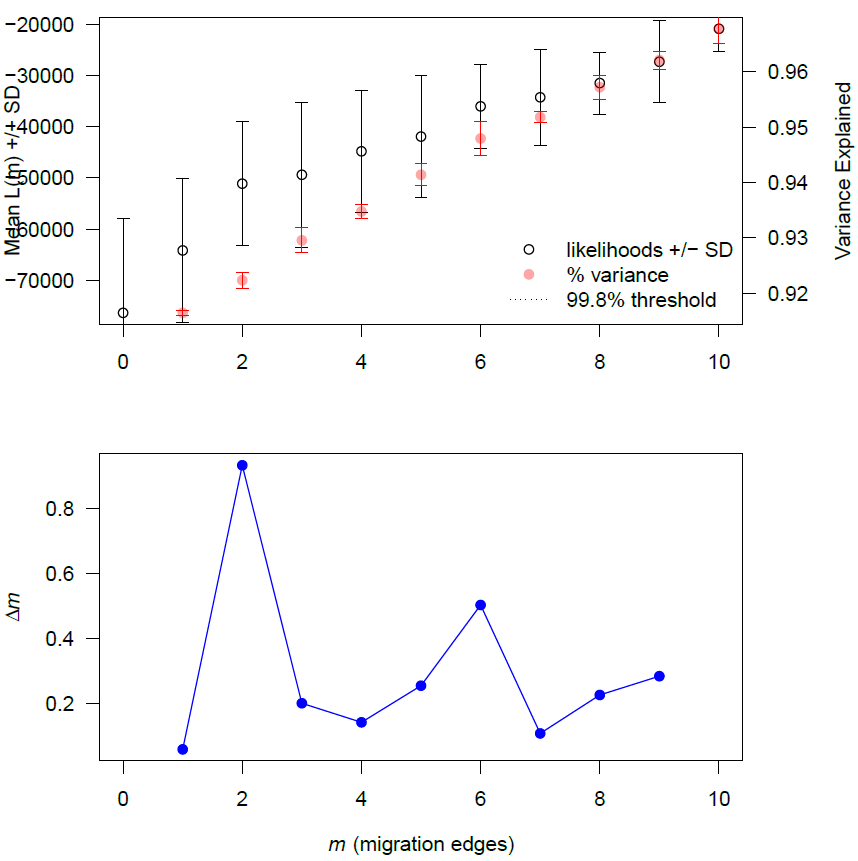
**Figure S2.** Evanno test of possible migration events between 27 naturally occurring populations, eastern wild Mediterranean population and cultivated accessions of olives using TreeMix. One hundred TreeMix runs were done with a random SNPs block size between 100 and 1000, from 1 to 10 migrations each with M29 as outgroup. The analysis was performed with *OptM* package.

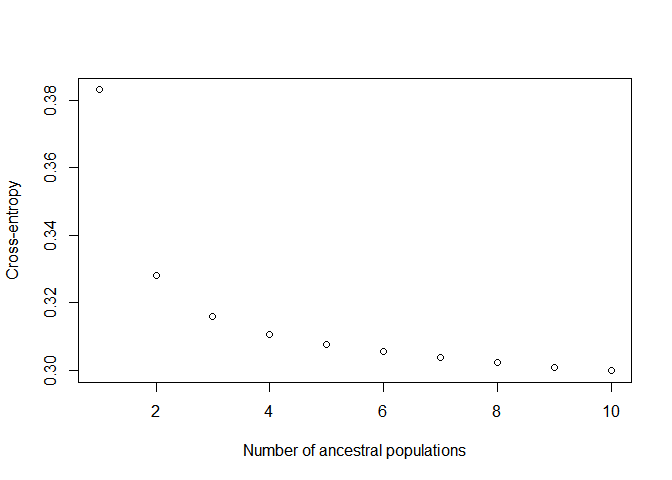
**Figure S3.** Cross-entropy criterion inferred by sNMF analyze performed on the genome-wide SNPs diversity of natural populations of *O. europaea L.* collected in France (143), Spain (123), Morocco (96) and Turkey (13) and cultivated *O. europaea L.* from the western Mediterranean Basin (145), using 142,060 SNPs.

**Table S1.** List of the baits used in the study. The baits were designed based on annotated gene of *Olea europaea* var. *europaea* (cv. Farga) Oe9 genome assembly (Julca et al. 2020). This table is in the supplementary xls file.

**Table S2.** Summary information of the sequenced cultivated accessions of *O. europaea* L. used in the study

| **Sample ID** | **Code** | **Accession origine** | | **Cultivar origine** | **Source** |
| --- | --- | --- | --- | --- | --- |
| Unkown-VS2-545 | MAR00545 | | Morocco | Morocco | World Olive Germplasm Bank of Marrakech |
| Acebuchera | MAR00215 | | Spain | Spain | World Olive Germplasm Bank of Marrakech |
| Aglandau | MAR00187 | | France | France | World Olive Germplasm Bank of Marrakech |
| Alameno Blanco | MAR00216 | | Spain | Spain | World Olive Germplasm Bank of Marrakech |
| Alameno de Montilla | MAR00218 | | Spain | Spain | World Olive Germplasm Bank of Marrakech |
| Amargoso | MAR00219 | | Spain | Spain | World Olive Germplasm Bank of Marrakech |
| Arbequina | MAR00220 | | Spain | Spain | World Olive Germplasm Bank of Marrakech |
| Azul | MAR00221 | | Spain | Spain | World Olive Germplasm Bank of Marrakech |
| Berri Meslal-397 | MAR00397 | | Morocco | Morocco | World Olive Germplasm Bank of Marrakech |
| Berri Meslal-532 | MAR00532 | | Morocco | Morocco | World Olive Germplasm Bank of Marrakech |
| Bical | MAR00333 | | Spain | Spain | World Olive Germplasm Bank of Marrakech |
| Blanqueta | MAR00222 | | Spain | Spain | World Olive Germplasm Bank of Marrakech |
| Bolvino | MAR00223 | | Spain | Spain | World Olive Germplasm Bank of Marrakech |
| Borriolenca | MAR00334 | | Spain | Spain | World Olive Germplasm Bank of Marrakech |
| Bouchouika | MAR00394 | | Morocco | Morocco | World Olive Germplasm Bank of Marrakech |
| Bouteillan | MAR00189 | | France | France | World Olive Germplasm Bank of Marrakech |
| Callosina | MAR00391 | | Morocco | Spain | World Olive Germplasm Bank of Marrakech |
| Canivano Negro | MAR00224 | | Spain | Spain | World Olive Germplasm Bank of Marrakech |
| Carrasqueno de Jumilla | MAR00226 | | Spain | Spain | World Olive Germplasm Bank of Marrakech |
| Carrasquillo | MAR00335 | | Spain | Spain | World Olive Germplasm Bank of Marrakech |
| Cayon | MAR00191 | | France | France | World Olive Germplasm Bank of Marrakech |
| Cerezuela | MAR00349 | | Spain | Spain | World Olive Germplasm Bank of Marrakech |
| Changlot Real | MAR00227 | | Spain | Spain | World Olive Germplasm Bank of Marrakech |
| Chorruo | MAR00229 | | Spain | Spain | World Olive Germplasm Bank of Marrakech |
| Cirujal | MAR0043 | | Italy | Spain | World Olive Germplasm Bank of Marrakech |
| Corbella | MAR00230 | | Spain | Spain | World Olive Germplasm Bank of Marrakech |
| Cornezuelo de Jaen | MAR00231 | | Spain | Spain | World Olive Germplasm Bank of Marrakech |
| Cornicabra | MAR00232 | | Spain | Spain | World Olive Germplasm Bank of Marrakech |
| Cucca | MAR0032 | | Spain | Spain | World Olive Germplasm Bank of Marrakech |
| Dolce di Rossano | MAR0011 | | Spain | Spain | World Olive Germplasm Bank of Marrakech |
| Dressi | MAR00286 | | Spain | Spain | World Olive Germplasm Bank of Marrakech |
| Dulzal | MAR00233 | | Spain | Spain | World Olive Germplasm Bank of Marrakech |
| El Lewa | MAR00494 | | Spain | Spain | World Olive Germplasm Bank of Marrakech |
| Empeltre | MAR00234 | | Spain | Spain | World Olive Germplasm Bank of Marrakech |
| Enagua de Arenas | MAR00336 | | Spain | Spain | World Olive Germplasm Bank of Marrakech |
| Escarabajuelo de Posadas | MAR00235 | | Spain | Spain | World Olive Germplasm Bank of Marrakech |
| Escarabajuelo de Úbeda | MAR00236 | | Spain | Spain | World Olive Germplasm Bank of Marrakech |
| Farga | MAR00338 | | Spain | Spain | World Olive Germplasm Bank of Marrakech |
| Frantoio | MAR0039 | | Spain | Spain | World Olive Germplasm Bank of Marrakech |
| Fulla de Salze | MAR00339 | | Spain | Spain | World Olive Germplasm Bank of Marrakech |
| Gentile di chieti | MAR0015 | | Spain | Spain | World Olive Germplasm Bank of Marrakech |
| Gordal Sevillana | MAR00108 | | Italy | Spain | World Olive Germplasm Bank of Marrakech |
| Gordal de Granada | MAR00238 | | Spain | Spain | World Olive Germplasm Bank of Marrakech |
| Grappolo | MAR0041 | | Spain | Spain | World Olive Germplasm Bank of Marrakech |
| Grossane-194 | MAR00194 | | France | France | World Olive Germplasm Bank of Marrakech |
| Habichuelero de Grazalema | MAR00239 | | Spain | Spain | World Olive Germplasm Bank of Marrakech |
| Hojiblanca | MAR00240 | | Spain | Spain | World Olive Germplasm Bank of Marrakech |
| Idleb | MAR00591 | | Spain | Spain | World Olive Germplasm Bank of Marrakech |
| Jabaluna | MAR00241 | | Spain | Spain | World Olive Germplasm Bank of Marrakech |
| Jaropo | MAR00242 | | Spain | Spain | World Olive Germplasm Bank of Marrakech |
| Khashabi-631 | MAR00631 | | Spain | Spain | World Olive Germplasm Bank of Marrakech |
| Lastrino | MAR0023 | | Spain | Spain | World Olive Germplasm Bank of Marrakech |
| Lazzero di prata | MAR00367 | | Spain | Spain | World Olive Germplasm Bank of Marrakech |
| Lechin de Sevilla | MAR00243 | | Spain | Spain | World Olive Germplasm Bank of Marrakech |
| Lechin de Granada | MAR00340 | | Spain | Spain | World Olive Germplasm Bank of Marrakech |
| Lentisca-244 | MAR00244 | | Spain | Spain | World Olive Germplasm Bank of Marrakech |
| Limoncillo | MAR00341 | | Spain | Spain | World Olive Germplasm Bank of Marrakech |
| Lloron de Atarfe | MAR00245 | | Spain | Spain | World Olive Germplasm Bank of Marrakech |
| Llumeta | MAR00343 | | Spain | Spain | World Olive Germplasm Bank of Marrakech |
| Loaime | MAR00344 | | Spain | Spain | World Olive Germplasm Bank of Marrakech |
| Lucques | MAR00195 | | France | France | World Olive Germplasm Bank of Marrakech |
| Machorron | MAR00247 | | Spain | Spain | World Olive Germplasm Bank of Marrakech |
| Manzanilla Cacerena | MAR00248 | | Spain | Spain | World Olive Germplasm Bank of Marrakech |
| Manzanilla de Sevilla | MAR00251 | | Spain | Spain | World Olive Germplasm Bank of Marrakech |
| Manzanilla de Agua | MAR00345 | | Spain | Spain | World Olive Germplasm Bank of Marrakech |
| Manzanilla de Hellin | MAR00346 | | Spain | Spain | World Olive Germplasm Bank of Marrakech |
| Manzanilla de Montefrio | MAR00250 | | Spain | Spain | World Olive Germplasm Bank of Marrakech |
| Mesyaf-641 | MAR00641 | | Spain | Spain | World Olive Germplasm Bank of Marrakech |
| Mignolo Cerretano | MAR0046 | | Spain | Spain | World Olive Germplasm Bank of Marrakech |
| Minekiri | MAR00634 | | Spain | Spain | World Olive Germplasm Bank of Marrakech |
| Mollar de Cieza | MAR00348 | | Spain | Spain | World Olive Germplasm Bank of Marrakech |
| Morchione | MAR0067 | | France | France | World Olive Germplasm Bank of Marrakech |
| Morisca | MAR00254 | | Spain | Morocco | World Olive Germplasm Bank of Marrakech |
| Morona | MAR00246 | | Spain | Spain | World Olive Germplasm Bank of Marrakech |
| Morrut | MAR00350 | | Spain | Spain | World Olive Germplasm Bank of Marrakech |
| Negral de Sabinan-255 | MAR00255 | | Spain | Spain | World Olive Germplasm Bank of Marrakech |
| Negrillo Redondo | MAR00257 | | Spain | Spain | World Olive Germplasm Bank of Marrakech |
| Negrillo de Arjona | MAR00256 | | Spain | Spain | World Olive Germplasm Bank of Marrakech |
| Negrillo de Estepa | MAR00351 | | Spain | Spain | World Olive Germplasm Bank of Marrakech |
| Negrillo de Iznalloz | MAR00352 | | Spain | Spain | World Olive Germplasm Bank of Marrakech |
| Nerba | MAR00123 | | Spain | Spain | World Olive Germplasm Bank of Marrakech |
| Nevado Azul | MAR00354 | | Spain | Spain | World Olive Germplasm Bank of Marrakech |
| Nevado Basto | MAR00225 | | Spain | Spain | World Olive Germplasm Bank of Marrakech |
| Nevado Rizado | MAR00355 | | Spain | Spain | World Olive Germplasm Bank of Marrakech |
| Ocal | MAR00258 | | Spain | Spain | World Olive Germplasm Bank of Marrakech |
| Ojo de Liebre | MAR00259 | | Spain | Spain | World Olive Germplasm Bank of Marrakech |
| Olivo de Mancha Real | MAR00260 | | Spain | Spain | World Olive Germplasm Bank of Marrakech |
| Olivo di Mandanici | MAR00136 | | Syria | Spain | World Olive Germplasm Bank of Marrakech |
| Palomar | MAR00262 | | Spain | Spain | World Olive Germplasm Bank of Marrakech |
| Patronet | MAR00263 | | Spain | Spain | World Olive Germplasm Bank of Marrakech |
| Picholine | MAR00196 | | France | France | World Olive Germplasm Bank of Marrakech |
| Picholine Marocaine | MAR00540 | | Morocco | Morocco | World Olive Germplasm Bank of Marrakech |
| Pico Limon de Grazalema | MAR00265 | | Spain | Spain | World Olive Germplasm Bank of Marrakech |
| Picual | MAR00267 | | Spain | Spain | World Olive Germplasm Bank of Marrakech |
| Picudo | MAR00356 | | Spain | Spain | World Olive Germplasm Bank of Marrakech |
| Plementa Bjelica | MAR00402 | | Spain | Spain | World Olive Germplasm Bank of Marrakech |
| Puntoza | MAR00499 | | Spain | Spain | World Olive Germplasm Bank of Marrakech |
| Racimal | MAR00268 | | Spain | Spain | World Olive Germplasm Bank of Marrakech |
| Rapasayo | MAR00357 | | Spain | Spain | World Olive Germplasm Bank of Marrakech |
| Razzaio | MAR0063 | | France | France | World Olive Germplasm Bank of Marrakech |
| Rechino | MAR00269 | | Spain | Spain | World Olive Germplasm Bank of Marrakech |
| Ronde de la Menara | MAR00543 | | Morocco | Morocco | World Olive Germplasm Bank of Marrakech |
| Rossellino | MAR0060 | | Spain | Spain | World Olive Germplasm Bank of Marrakech |
| Royal de Cazorla | MAR00270 | | Spain | Spain | World Olive Germplasm Bank of Marrakech |
| Sabatera | MAR00271 | | Spain | Spain | World Olive Germplasm Bank of Marrakech |
| Salonenque | MAR00197 | | France | France | World Olive Germplasm Bank of Marrakech |
| Santa Martinenga | MAR00145 | | Syria | Spain | World Olive Germplasm Bank of Marrakech |
| Sayali | MAR00287 | | Spain | Spain | World Olive Germplasm Bank of Marrakech |
| Sevillano de Jumilla | MAR00272 | | Spain | Spain | World Olive Germplasm Bank of Marrakech |
| Sevillenca | MAR00358 | | Spain | Spain | World Olive Germplasm Bank of Marrakech |
| Sinopolese | MAR0018 | | Spain | Spain | World Olive Germplasm Bank of Marrakech |
| Storta | MAR00406 | | Spain | Spain | World Olive Germplasm Bank of Marrakech |
| Tabelout | MAR00437 | | Spain | Spain | World Olive Germplasm Bank of Marrakech |
| Tebabs | MAR00661 | | Spain | Spain | World Olive Germplasm Bank of Marrakech |
| Teffah | MAR00427 | | Spain | Spain | World Olive Germplasm Bank of Marrakech |
| Tempranillo de Yeste-274 | MAR00274 | | Spain | Spain | World Olive Germplasm Bank of Marrakech |
| Tonda Iblea | MAR0012 | | Spain | Spain | World Olive Germplasm Bank of Marrakech |
| Unkown-OT2-537 | MAR00537 | | Morocco | Morocco | World Olive Germplasm Bank of Marrakech |
| Unkown-VS1-544 | MAR00544 | | Spain | Spain | World Olive Germplasm Bank of Marrakech |
| Unkown-VS2-545 | MAR00546 | | Morocco | Morocco | World Olive Germplasm Bank of Marrakech |
| Unkown-VS2-545 | MAR00545 | | Morocco | Morocco | World Olive Germplasm Bank of Marrakech |
| Unkown-VS5-547 | MAR00547 | | Morocco | Morocco | World Olive Germplasm Bank of Marrakech |
| Uovo di Piccione | MAR00141 | | Syria | Spain | World Olive Germplasm Bank of Marrakech |
| Varudo | MAR00273 | | Spain | Spain | World Olive Germplasm Bank of Marrakech |
| Varudo-275 | MAR00275 | | Spain | Spain | World Olive Germplasm Bank of Marrakech |
| Vera | MAR00276 | | Spain | Spain | World Olive Germplasm Bank of Marrakech |
| Verdala | MAR00278 | | Spain | Spain | World Olive Germplasm Bank of Marrakech |
| Verdale | MAR00199 | | France | France | World Olive Germplasm Bank of Marrakech |
| Verdial de Badajoz | MAR00342 | | Spain | Spain | World Olive Germplasm Bank of Marrakech |
| Verdial de Huevar | MAR00213 | | Portugal | Spain | World Olive Germplasm Bank of Marrakech |
| Verdiell | MAR00279 | | Spain | Spain | World Olive Germplasm Bank of Marrakech |
| Villalonga | MAR00201 | | Portugal | Spain | World Olive Germplasm Bank of Marrakech |
| Zalmati-299 | MAR00299 | | Spain | Spain | World Olive Germplasm Bank of Marrakech |
| Zarza | MAR00280 | | Spain | Spain | World Olive Germplasm Bank of Marrakech |
| Zeletni | MAR00421 | | Spain | Spain | World Olive Germplasm Bank of Marrakech |
| Zinzala | CTO_E8_N4 | | France | France | Technical Centrer of Olive |
| Petit Ribier | CTO_P10_F06 | | France | France | Technical Centrer of Olive |
| Sabine | CTO_P10_N27 | | France | France | Technical Centrer of Olive |
| Cayet Roux | POR_P39_33_10 | | France | France | Porquerolles collection |
| Salonenque | POR_P39_33_16 | | France | France | Porquerolles collection |
| Brun | POR_P39_37_21 | | France | France | Porquerolles collection |
| Tanche | POR_P43_02_05 | | France | France | Porquerolles collection |
| Cayon | POR_P43_03_12 | | France | France | Porquerolles collection |
| Reymet | POR_P43_06_01 | | France | France | Porquerolles collection |
| Grossane | POR_P43_17_13 | | France | France | Porquerolles collection |

**Table S3.** List of filters and remaining SNPs for each on genomic data of target sequencing of 561 genotypes of *O. europaea* L.

|  | **Total SNPs remaining** |
| --- | --- |
| **Raw data** | 27275679 |
| **Remove Indel** | 24547413 |
| **Quality > 200** | 22340559 |
| **SNP Biallelic only** | 21036126 |
| **Min mean depth 8** | 958648 |
| **Max mean depth 400** | 957569 |
| **SNP cluster (3,10)** | 383904 |
| **Min depth 8** | 383904 |
| **Missing data Site > 0.15** | 155063 |
| **heterozygoty > 0.85** | 154675 |
| **Minor allele count 1** | 142060 |

**Table S4.** List of individuals of *O. europaea* L. removed from genomic data set because of missing data >0.2

| **Sample ID removed** | **Type** |
| --- | --- |
| Manzanilla_de_Abla | Cultivated |
| OES_E13_03 | Wild |
| OES_E15_08 | Wild |
| OES_E15_12 | Wild |
| OES_E16_05 | Wild |
| OES_E16_10 | Wild |
| OES_E16_13 | Wild |
| OES_E17_08 | Wild |
| OES_E18_02 | Wild |
| OES_E18_04 | Wild |
| OES_E18_11 | Wild |
| OES_E19_12 | Wild |
| OES_E20_02 | Wild |
| OES_F02_01 | Wild |
| OES_F02_15 | Wild |
| OES_F07_08 | Wild |
| OES_F07_10 | Wild |
| OES_F10_05 | Wild |
| OES_F10_13 | Wild |
| OES_F11_04 | Wild |
| OES_M21_03 | Wild |
| OES_M21_13 | Wild |
| OES_M23_03 | Wild |
| OES_M23_05 | Wild |
| OES_M23_07 | Wild |
| OES_M23_09 | Wild |
| OES_M23_10 | Wild |
| OES_M23_11 | Wild |
| OES_M23_13 | Wild |
| OES_M23_14 | Wild |
| OES_M23_15 | Wild |
| OES_M28_10 | Wild |
| OES_M29_04 | Wild |
| OST072 | Wild |
| OST082 | Wild |

**Table S5.** Results of bootstraping over loci of pairwise *F*_ST_ performed on the genome-wide SNPs diversity of natural populations of *O. europaea* L. collected in France (143), Spain (123), Morocco (96) and Turkey (13) using 142,060 SNPs. The upper limit ci is the upper part and the lower limit ci in the bottom part of the matrix. This table is in the supplementary xls file.

**Table S6**. List of cultivated accessions of *O. europaea* L. and their assignment to genetic cluster (C0, C1, C3 or C4) according to sNMF assignation at K = 4. The detailed method is developed above (Supplementary information)

| **Accession names** | **Cluster assignation** |
| --- | --- |
| Unkown-VS2-545 | C0 |
| Acebuchera | C4 |
| Aglandau | C1 |
| Alameno Blanco | C4 |
| Alameno de Montilla | C0 |
| Amargoso | C1 |
| Arbequina | C3 |
| Azul | C0 |
| Berri Meslal-397 | C0 |
| Berri Meslal-532 | C0 |
| Bical | C4 |
| Blanqueta | C3 |
| Bolvino | C0 |
| Borriolenca | C0 |
| Bouchouika | C4 |
| Bouteillan | C1 |
| Zinzala | C3 |
| Petit Ribier | C3 |
| Sabine | C0 |
| Callosina | C4 |
| Canivano Negro | C4 |
| Carrasqueno de Jumilla | C4 |
| Carrasquillo | C4 |
| Cayon | C1 |
| Cerezuela | C4 |
| Changlot Real | C0 |
| Chorruo | C4 |
| Cirujal | C0 |
| Corbella | C0 |
| Cornezuelo de Jaen | C0 |
| Cornicabra | C4 |
| Cucca | C0 |
| Dolce di Rossano | C0 |
| Dressi | C0 |
| Dulzal | C4 |
| El Lewa | C1 |
| Empeltre | C0 |
| Enagua de Arenas | C4 |
| Escarabajuelo de Posadas | C4 |
| Escarabajuelo de Úbeda | C0 |
| Farga | C0 |
| Frantoio | C3 |
| Fulla de Salze | C4 |
| Gentile di chieti | C3 |
| Gordal Sevillana | C1 |
| Gordal de Granada | C4 |
| Grappolo | C0 |
| Grossane-194 | C1 |
| Habichuelero de Grazalema | C4 |
| Hojiblanca | C4 |
| Idleb | C1 |
| Jabaluna | C0 |
| Jaropo | C4 |
| Khashabi-631 | C1 |
| Lastrino | C0 |
| Lazzero di prata | C3 |
| Lechin de Sevilla | C0 |
| Lechin de Granada | C0 |
| Lentisca-244 | C0 |
| Limoncillo | C4 |
| Lloron de Atarfe | C4 |
| Llumeta | C0 |
| Loaime | C4 |
| Lucques | C1 |
| Machorron | C4 |
| Manzanilla Cacerena | C4 |
| Manzanilla de Sevilla | C4 |
| Manzanilla de Agua | C4 |
| Manzanilla de Hellin | C4 |
| Manzanilla de Montefrio | C4 |
| Mesyaf-641 | C1 |
| Mignolo Cerretano | C3 |
| Minekiri | C1 |
| Mollar de Cieza | C4 |
| Morchione | C0 |
| Morisca | C4 |
| Morona | C4 |
| Morrut | C0 |
| Negral de Sabinan-255 | C1 |
| Negrillo Redondo | C4 |
| Negrillo de Arjona | C4 |
| Negrillo de Estepa | C4 |
| Negrillo de Iznalloz | C4 |
| Nerba | C1 |
| Nevado Azul | C4 |
| Nevado Basto | C4 |
| Nevado Rizado | C4 |
| Ocal | C4 |
| Ojo de Liebre | C4 |
| Olivo de Mancha Real | C4 |
| Olivo di Mandanici | C3 |
| Cayet Roux | C0 |
| Salonenque | C1 |
| Brun | C1 |
| Tanche | C1 |
| Cayon | C0 |
| Reymet | C3 |
| Grossane | C1 |
| Palomar | C0 |
| Patronet | C0 |
| Picholine | C0 |
| Picholine Marocaine | C4 |
| Pico Limon de Grazalema | C4 |
| Picual | C4 |
| Picudo | C4 |
| Plementa Bjelica | C0 |
| Puntoza | C0 |
| Racimal | C4 |
| Rapasayo | C0 |
| Razzaio | C3 |
| Rechino | C4 |
| Ronde de la Menara | C4 |
| Rossellino | C3 |
| Royal de Cazorla | C4 |
| Sabatera | C0 |
| Salonenque | C1 |
| Santa Martinenga | C0 |
| Sayali | C0 |
| Sevillano de Jumilla | C4 |
| Sevillenca | C0 |
| Sinopolese | C0 |
| Storta | C0 |
| Tabelout | C3 |
| Tebabs | C1 |
| Teffah | C1 |
| Tempranillo de Yeste-274 | C4 |
| Tonda Iblea | C1 |
| Unkown-OT2-537 | C3 |
| Unkown-VS1-544 | C0 |
| Unkown-VS2-545 | C0 |
| Unkown-VS2-545 | C0 |
| Unkown-VS5-547 | C0 |
| Uovo di Piccione | C1 |
| Varudo | C4 |
| Varudo-275 | C4 |
| Vera | C3 |
| Verdala | C4 |
| Verdale | C1 |
| Verdial de Badajoz | C4 |
| Verdial de Huevar | C4 |
| Verdiell | C0 |
| Villalonga | C0 |
| Zalmati-299 | C3 |
| Zarza | C0 |
| Zeletni | C0 |

Table S7. Descriptive data of depth and enrichment rate of *O. europaea* L. using target sequencing method versus whole genome sequencing method.

| **Sample ID** | **Mean depth whole genome** | **Mean depth target sequencing** | **Enrichment rate** |
| --- | --- | --- | --- |
| OES_E13_09 | 1.21 | 40.40 | 33.27 |
| OES_F10_03 | 1.27 | 34.72 | 27.31 |
| Picholine | 0.77 | 18.20 | 23.59 |
| Picholine_Marocaine | 1.27 | 67.20 | 52.88 |
| Mean_all | 1.13 | 40.13 | 34.26 |
